# Astroglial dysfunctions drive aberrant synaptogenesis in developing brain with lengthy general anesthesia

**DOI:** 10.1101/477075

**Authors:** Bin Zhou, Lingmin Chen, Ping Liao, Lu Huang, Zhuo Chen, Guoqiang Yu, Li Wang, Jianguo Zhang, Yunxia Zuo, Jin Liu, Ruotian Jiang

## Abstract

Lengthy use of general anesthetics (GAs) causes cognitive deficits in developing brain, which has raised significant clinical concerns such that FDA is warning on the use of GAs in children younger than 3 years. However, the molecular and cellular mechanisms for GAs-induced neurotoxicity remain largely unknown. Here we report that sevoflurane, a commonly used GA in pediatrics, causes compromised astrocyte morphogenesis, spatiotemporally correlated to the synaptic overgrowth with reduced synaptic function in developing cortex in a regional-, exposure-length- and age-specific manner. Sevoflurane disrupts astrocyte Ca^2+^ homeostasis both acutely and chronically, which leads to the down regulation of Ezrin, an actin-binding membrane protein, which we found is critically involved in astrocyte morphogenesis *in vivo*. Importantly, in normal developing brain, the genetic intervention of astrocyte morphogenesis is sufficient to produce the aberrant synaptic structure and function virtually identical to the ones induced by lengthy sevoflurane exposure. Our data uncover that astrocytes are unexpectedly central targets for GAs to exert toxic effects, and that astrocyte morphological integrity is crucial for synaptogenesis in the developing brain.

## Introduction

Millions infants and young children are undergoing general anesthesia every year. Human retrospective cohort studies strongly suggest that exposure of GAs and sedation drugs to immature individuals produced developmental and behavioral disorders and learning disabilities (1, 2). The anesthetic-induced developmental neurotoxicity (AIDN) has raised significant concerns to clinicians, scientists and the public about the safety of general anesthesia to infants and young children. In the light of the current clinical and preclinical evidence, in 2016, the U.S. Food and Drug Administration (FDA) has issued a drug safety communication warning that “*repeated or lengthy use of general anesthetic and sedation drugs in children younger than 3 years or in pregnant women may affect the development of children’s brains*.” (https://www.fda.gov/Drugs/DrugSafety/ucm532356.htm). Importantly, two recent cohort studies (3, 4) further suggest that multiple (but not a single) GAs exposure, is associated with decreases in processing speed and fine motor coordination. In summary, the existing clinical evidence is sufficiently concerning that more efforts are required to elucidate the mechanisms by which GAs exposure, particularly lengthy or repeat exposure, affects brain development.

Preclinical studies have clearly shown that early lengthy/repeated exposure of nearly all commonly used GAs, including sevoflurane, isoflurane, propofol, ketamine, cause cognitive dysfunctions in the developing rodent brain (1, 2). These disorders include spatial (2), no-spatial (5) and fear condition learning deficits (6) in rodents, anxiety-related behaviors or motor reflex deficits in non-human primates(1). However, the underlying molecular and cellular mechanisms are not completely understood. Early studies mainly focused on apoptotic hypothesis. Nevertheless, in most studies, only a very small fraction number of cells displayed activated caspase-3 expression after GAs exposure, and there is no strong evidence for a causal link between neuronal apoptosis and lasting neurocognitive impairment (1). Recently, it has been shown that early lengthy exposure of GAs induced increased or decreased dendrite outgrowth, disruption of axon guidance, and altered synaptic functions in mouse cortex and hippocampus (1, 5, 7, 8). Aberrant synaptogenesis and neural circuit formation have strong causal links with cognitive dysfunctions, which has been clearly demonstrated in many psychiatric and neurodevelopmental disorders. However, little is known regarding the molecular and cellular mechanisms underlying the synaptic and circuit dysfunctions induced by GAs during the critical period of the brain development.

Astrocytes constitute at least one-third of human brain cells. Astrocyte processes contact with axons and dendritic spines, forming “tripartite synapses”, and play vital roles in the process of synaptogenesis during brain development (9, 10). Sparse studies from the literature suggest that lengthy GAs exposure altered astrocyte functions *in vitro*, including impaired expressions of glial fibrillary acidic protein (GFAP), glutamate-aspartate transporter (GLAST) and brain-derived neurotrophic factor (BDNF) (11, 12), and astrocyte capability to support neuronal development (13). However, most of the studies described above either used *in vitro* exposure to cultured astrocytes, which differ significantly from general anesthesia *in vivo*, or provided little information on how astrocyte dysfunctions correlated or contributed to measurable neuronal deficits under the same experimental settings. Hence, how GAs affect astrocyte developmental biology and how GAs disrupt glia-neuron interactions during synaptogenesis *in vivo* remain largely unexplored.

In the current study, in combination of Ca^2+^ imaging, high-resolution morphological reconstructions using light and electron microscopy and mouse genetics, we explored in detail how sevoflurane (Sevo), a commonly used general anesthetic in pediatrics, impacts negatively astrocyte morphogenesis and Ca^2+^ signaling *in situ* and *in vivo*, and provided evidence to show that astrocyte dysfunctions drive neuronal dysfunctions in the developing somatosensory cortex with lengthy Sevo exposure.

## Results

### Lengthy Sevo exposure disrupts astrocyte morphogenesis with compromised “tripartite synapse” maturation in developing somatosensory cortex

To evaluate how lengthy Sevo exposure affects developing mouse brain, we had P7 mice exposed to 2.5% Sevo (we termed hereafter Sevo group mice) or their littermates exposed to carrier gas (30% O_2_/70% CO_2_) (we termed hereafter Control group mice) for 4 h. This experimental setting was chosen on the basis of a number of clinical studies which have shown that cognitive dysfunctions were observed in children/infants with GAs exposure longer than 3 h (14). All mice survived after exposure, with both artery blood gas parameters measured immediately after exposure **(Supplementary Table1)** and body weight at P14 comparable to the Control group mice (Control group vs Sevo group: 6.55 ± 0.73 g vs 6.27 ± 0.73 g, *P* = 0.299, n = 16 mice per group, unpaired *t* test).

Then, we examined in detail how lengthy Sevo exposure affected astrocyte morphogenesis during the critical period of the brain development, i.e., from P7 to P21 in mice (10). To achieve this, we performed intracellular lucifer yellow iontophoresis in lightly fixed tissue followed with confocal imaging and morphological 3D reconstructions to analyze astrocyte morphology *in situ* (15) at P8, P14 and P21 (**Fig. 1A**). In somatosensory cortex, astrocytes from Sevo group mice displayed significantly smaller distal (away from soma) and proximal (closed to soma) fine process volume, with decreased territory volume, measured at both P8 and P14. However, when at P21, there was no more significance between the two groups (**Fig. 1B, C**). No significant difference was observed between the two groups in astrocyte soma and primary branches (branches directly protruding from soma) at P8, P14 and P21 (**Supplementary Fig. 1A, B**). We also quantified GFAP expressions using immunostaining, a well-known marker for astrogliosis, and no significant difference between the two groups was found (**Supplementary Fig. 1C, D**). Therefore, our data so far suggest that Sevo exposure induced compromised astrocyte fine structure development without causing apparent astrogliosis in the mouse cortex.

**Fig 1.**
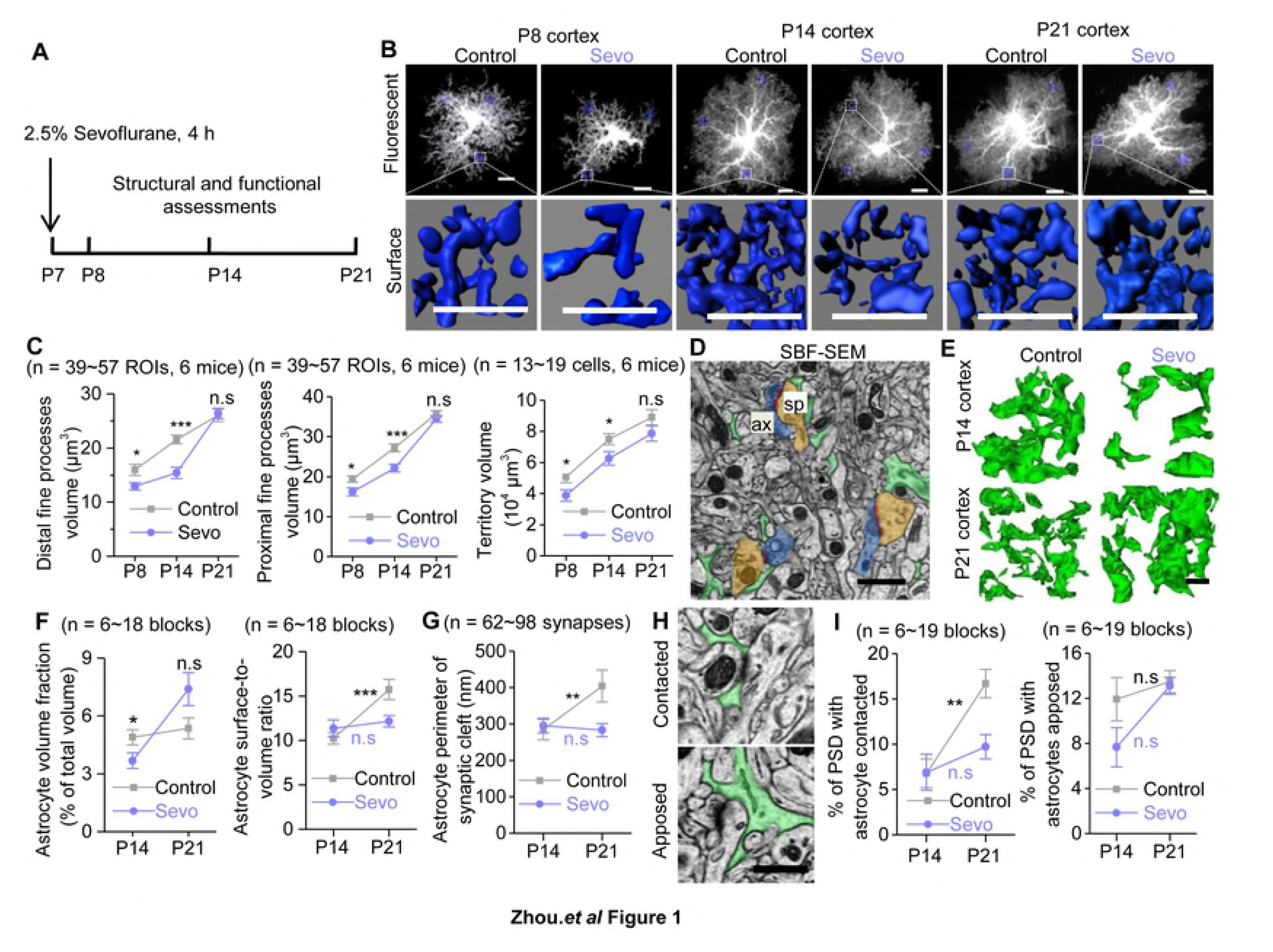
Lengthy Sevo exposure disrupts astrocyte morphogenesis and compromised “tripartite synapse” maturation in developing somatosensory cortex. (**A**) Schematic illustrating the experimental protocol for Sevo exposure. (**B**) Representative confocal images and 3D reconstructed distal fine processes of sparsely labeled astrocytes from Control and Sevo group mice at P8, P14 and P21. Scale bars, 10 μm. (**C**) **Left:** Quantification of astrocyte distal fine processes volume in Control and Sevo group mice at P8 (*P =* 0.02, unpaired *t* test), P14 (*P* = 6.1E-6, unpaired *t* test) and P21 (*P* = 0.82, unpaired *t* test). **Middle:** Quantification of astrocyte proximal fine processes volume from Control and Sevo group mice at P8 (*P =* 0.012, unpaired *t* test), P14 (*P* = 3.1E-4, unpaired *t* test) and P21 (*P* = 0.279, unpaired *t* test). **Right:** Quantification of astrocyte territory volume from Control and Sevo group mice at P8 (*P =* 0.028, unpaired *t* test), P14 (*P* = 0.03, unpaired *t* test) and P21 (*P* = 0.248, unpaired *t* test). (**D**) SBF-SEM images of astrocyte fine processes (green), postsynaptic density (PSD, red), axonal bouton (ax, blue) and dendritic spine (sp, brown). Scale bar, 1 μm. (**E**) 3D reconstruction of astrocyte fine processes in Control and Sevo group mice at P14 and P21. Scale bar, 1 μm. (**F**) **Left:** Comparison of astrocyte volume fraction in Control and Sevo group mice at P14 (*P* = 0.038, unpaired *t* test) and P21 (*P* = 0.074, unpaired *t* test). **Right:** Increased astrocyte surface-to-volume ratio from P14 to P21 mice in Control (*P* = 3.6E-4, unpaired *t* test), but not in Sevo group mice (*P* = 0.628, unpaired *t* test). (**G**) Increased astrocyte perimeter enwrapping the synaptic cleft from P14 to P21 mice in Control (*P* = 0.007, Mann-Whitney test), but not in Sevo group mice (*P* = 0.232, Mann-Whitney test). (**H**) SBF-SEM images of astrocytes contacted or apposed PSD. Scale bar, 0.5 μm. (**I**) **Left:** Increased PSD with astrocytes contacted from P14 to P21 mice in Control (*P* = 0.002, Mann-Whitney test), but not in Sevo group mice (*P* = 0.129, Mann-Whitney test). **Right:** Unchanged PSD with astrocytes apposed from P14 to P21 mice in Control (*P* = 0.105, Mann-Whitney test) and Sevo (*P* = 0.07, unpaired *t* test) group. \**P* < 0.05, \*\**P* < 0.01, \*\*\**P* < 0.001, n.s, not significant. Data are shown as mean ± s.e.m.

In contrast to the cortex, much less morphological deficits were observed in the CA1 stratum radiatum (CA1sr) or in the molecular layer of dentate gyrus (DG-mo) of the hippocampus at P14 (**Supplementary Fig. 2**), a time point when cortical astrocytes showed remarkable morphological deficits in Sevo group mice. In clinical studies, cognitive dysfunctions are mainly associated with early childhood exposure to lengthy general anesthesia. We also tested if developing astrocytes were resistant to a relatively short Sevo exposure and if mature astrocytes were resistant to a 4 h Sevo exposure. Interestingly, neither cortical astrocytes in P7 mice with only 1 h Sevo exposure measured at P14 nor those in adult mice (~P45) with 4 h Sevo exposure measured 7 d later (**Supplementary Fig. 3**) displayed morphological deficits. Together, our data reveal that lengthy, but not short, Sevo exposure induces compromised astrocyte morphogenesis only in developing but not in mature brain.

Astrocyte processes are extremely fine (<50 nm) and form “tripartite synapse” with axonal boutons and dendritic spines (16). We then performed serial block face scanning electron microscopy (SBF-SEM) to examine the ultrastructure of the tripartite synapse. Lower astrocyte volume fraction (which reflects the absolute volume of astrocyte processes) was observed in the somatosensory cortex of Sevo group mice at P14, but not at P21. From P14 to P21, the surface-to-volume ratio (which reflects the fineness of astrocyte processes) was markedly increased by ~50% in Control group mice, but remained unchanged in Sevo group mice (**Fig. 1D-F**). Then we examined astrocyte-neuron contact. There was also an increase of astrocyte perimeters enwrapping the synaptic cleft by ~30% from P14 to P21 in Control group mice, but remained unchanged in Sevo group mice (**Fig. 1G**). In addition, there was also a more than one-fold increase of astrocyte fine processes insertion into synaptic cleft from P14 to P21, as shown by the markedly increased number of astrocyte contacted PSD in Control group mice but remained unchanged in Sevo group mice. The number of astrocyte apposed PSD remained unchanged in both groups from P14 to P21 (**Fig. 1H, I**).

The light and electron microscopic data together demonstrate that lengthy Sevo exposure disrupts astrocyte morphogenesis resulting in altered tripartite synaptic structure in a regional-, exposure-length- and age-specific manner.

### Aberrant synaptic growth and functions spatiotemporally correlated with astrocyte morphological deficits in Sevo group mice

Next, we tested whether lengthy Sevo exposure also leads to structural and functional deficits in cortical pyramidal neurons at P21. By performing lucifer yellow iontophersis in combine with morphological 3D reconstruction, we found that the pyramidal neurons in Sevo group mice had higher total and mushroom basal dendritic spine density than those in Control group mice at P21 (**Fig. 2A, B**). Interestingly, the dendritic spine density was similar in the hippocampal CA1sr (**Supplementary Fig. 4**) between the two groups, recalling the unchanged astrocyte morphology in this region. Using SBF-SEM, we identified structurally visible PSDs and reconstructed the corresponding dendritic spines as indictors for excitatory synapses (**Fig. 2C**). We found a more than 50% higher total dendritic spine density, whereas the mushroom spine density remained unchanged in the primary somatosensory cortex in Sevo group mice (**Fig. 2D**). This dataset suggests that the synaptic overgrowth was present in parallel with astrocytic structural deficits in the cortex but not in the hippocampus of Sevo group mice, and SBF-SEM data suggest the synaptic overgrowth largely resulted from an increase of synapses which are relatively immature or less functional.

**Fig 2.**
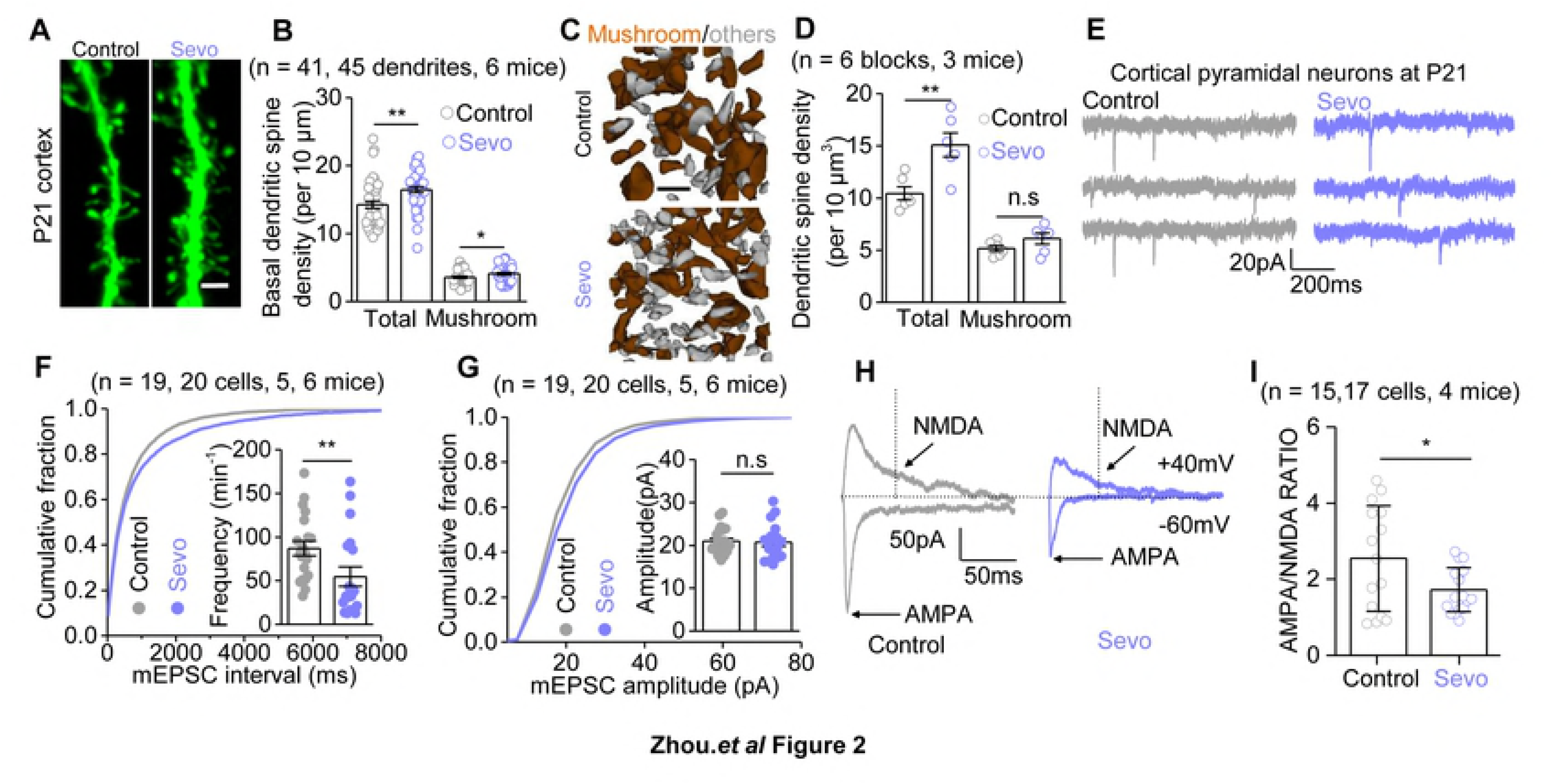
Aberrant synaptic growth and functions in Sevo group mice. (**A**) Representative images of basal dendritic spines in the somatosensory cortex of Control and Sevo group mice at P21. Scale bars, 2 μm. (**B**) Quantification of the total (*P* = 0.0016, unpaired *t* test) and mushroom (*P* = 0.039, unpaired *t* test) dendritic spine density in Control and Sevo group mice at P21. (**C**) 3D reconstruction of mushroom spines (brown) and other morphologically immature spines (grey). Scale bars, 1 μm. (**D**) Quantification of total (*P* = 0.005, unpaired *t* test) and mushroom (*P* = 0.155, unpaired *t* test) dendritic spine density in the somatosensory cortex of Control and Sevo group mice at P21 by using SBF-SEM. (**E**) Representative mEPSCs traces from L3-5 pyramidal neurons in the somatosensory cortex. **(F**) Cumulative distributions and average frequency (*P* = 0.007, Mann-Whitney test) of mEPSCs. (**G**) Cumulative distributions and average (*P* = 0.833, *t* test) of mEPSCs amplitude. (**H**) Representative eEPSCs traces from L3-5 pyramidal neurons from Control and Sevo mice at P21-P23. AMPAR-eEPSCs were recorded with the presence of 100 μM picrotoxinin and 20 μM glycine at −60 mV, and mixed AMPA and NMDA currents were recorded with 100 μM picrotoxinin and 20 μM glycine at +40 mV. The black dotted line (50 ms after the stimulation) indicates the measuring points for NMDAR-eEPSCs amplitudes. (**I**) Quantification of AMPA/NMDA ratio (*P* = 0.033, *t* test). \**P* < 0.05, \*\**P* < 0.01, \*\*\**P* < 0.001, n.s, not significant. Data are shown as mean ± s.e.m.

Was the synaptic overgrowth associated with enhanced synaptic function? To address this question, we measured AMPAR-mediated miniature EPSCs (mEPSCs) from pyramidal neurons in the cortical layer 3-5 using whole cell patch-clamp technique (**Fig. 2E**). Unexpectedly, the frequency of mEPSCs was not increased but instead markedly decreased by ~37% in the Sevo group mice, compared with the Control group mice (**Fig. 2F**), whereas the amplitudes of mEPSCs were similar **(Fig. 2G).** This dataset suggests that Sevo exposure resulted in a decrease of glutamate release probability or (and) a decrease of the number of AMPAR-containing synapses. Then we measured the evoked EPSCs (eEPSCs) in pyramidal neurons (**Fig. 2H**). We confirmed that the evoked currents were truly AMPA and NDMA mediated currents by using AMPA and NMDA specific antagonists CNQX and DL-AP5, respectively (**Supplementary Fig. 5A**). We found that the AMPAR/NMDAR ratio was significantly ~33% smaller in Sevo group when compared with Control group (**Fig. 2H, I**), with unchanged decay kinetics (**Supplementary Fig. 5B, C**). Since the reduction (37%) in mEPSC frequency can quantitatively account for the reduction in AMPAR/NMDAR ratio (33%) observed in Sevo group mice, these results suggest that the synaptic NMDAR function is largely unaffected in these mice. The structural and functional data together reveal that early Sevo exposure induces an increase of synapse number, some of which are very likely non-functional/immature, with reduced AMPA-mediated synaptic transmission in the cortex where astrocytes display compromised morphogenesis.

### Sevo exposure disrupted Ca^2+^ signals in developing cortical astrocytes

Next, we explored the mechanisms underlying the compromised astrocyte morphogenesis in Sevo group mice. Thrane *et al* reported that GAs (ketamine/xylazine, isoflurane, and urethane) exposure suppressed acutely spontaneous Ca^2+^ signals and evoked Ca^2+^ responses *in vivo* in mature astrocytes (17), which prompted us to measure the acute and chronic effects of lengthy Sevo exposure on developing cortical astrocytes. AAV5⋅gfaABC_1_D⋅GCaMP6f (GCaMP6f) was expressed in cortical astrocytes using *in vivo* microinjections at P0 (**Fig. 3A**) (**see methods**). GCaMP6f microinjections at P0 did not induce apparent astrogliosis (**Supplementary Fig. 6**) at P14.

**Fig 3.**
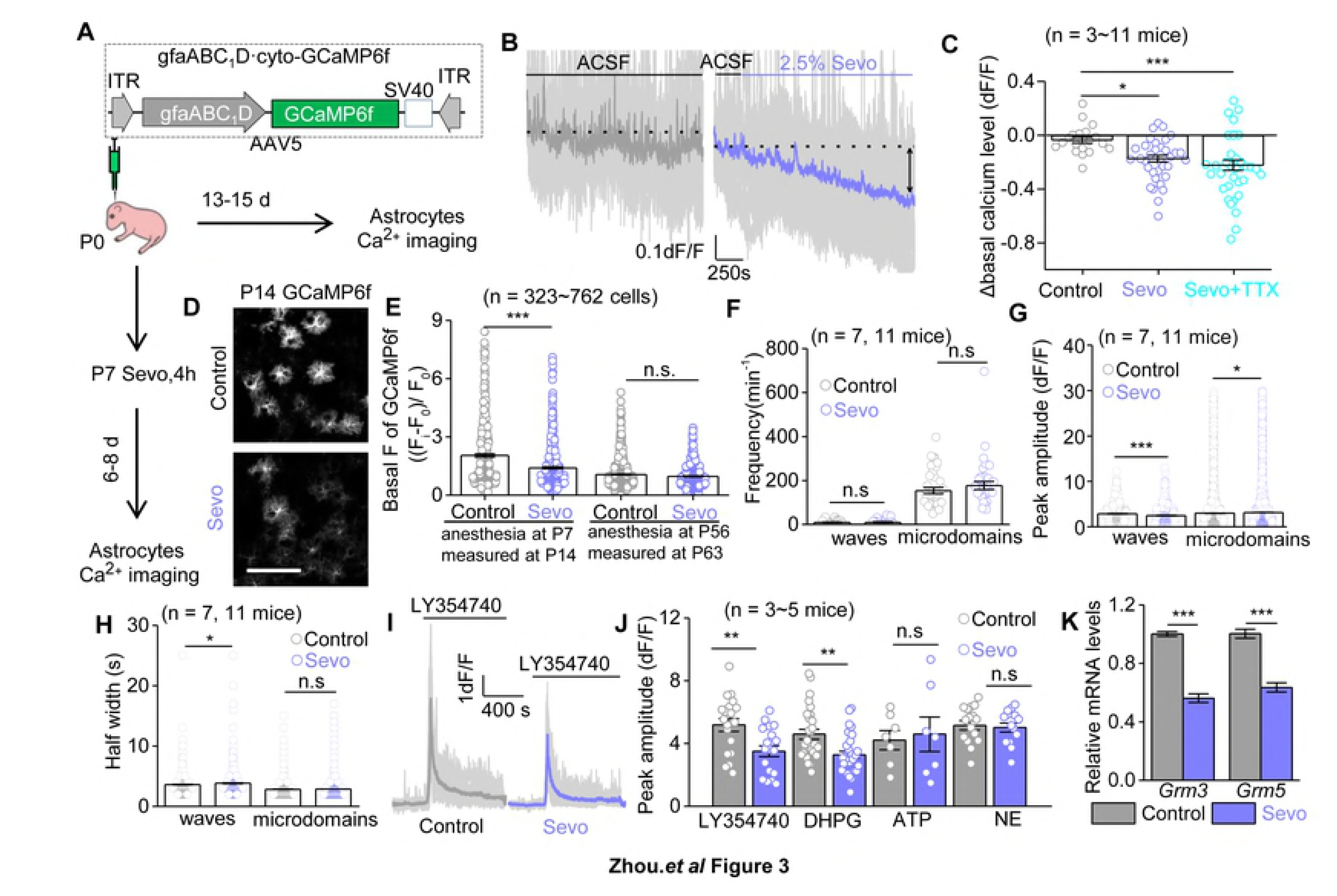
Sevo exposure disrupts Ca^2+^ signals in developing cortical astrocytes. (**A**) Schematic of Ca^2+^ imaging protocols. (**B**) Individual traces (light gray) and average (purple) trace showing altered astrocyte basal Ca^2+^ level by Sevo exposure. (**C**) Quantification of the effect of Sevo (*P* = 0.015, one-way ANOVA test; *post-hoc* Tukey test), Sevo + TTX (*P* = 6.8E-4, one-way ANOVA test; *post-hoc* Tukey test) on cortical astrocyte basal Ca^2+^ levels. (**D**) Representative confocal images of GCaMP6f-expressing astrocytes. Scale bar, 100 μm. (**E**) Quantification of basal fluorescent intensity of GCaMP6f (Control *vs* Sevo at P14, *P* = 6.4E-9, Mann-Whitney test, Control *vs* Sevo at P56, *P* = 0.523, Mann-Whitney test), F_i_: fluorescent intensity of GCaMP6f expressing cells; F_0:_ background intensity. (**F, G, H**) Quantification of frequency, peak amplitude and half-width for the two types of spontaneous Ca^2+^ signals at P13-P15. (*P* = 0.672, for waves frequency; *P* = 0.314, for microdomains frequency; *P* = 5.8E-5 for waves peak amplitude; *P* = 0.037 for microdomains peak amplitude; *P* = 0.012 for waves half-width; *P* = 0.771 for microdomains half-width, Mann-Whitney test). (**I**) Individual traces (light gray) and averages for Ca^2+^ responses to LY354770 (10 μM) at P13-P15. (**J**) Summary bar graph for peak amplitude of Ca^2+^ signals evoked by LY354740 (*P* = 0.004, unpaired *t* test), DHPG (*P* = 0.001, Mann-Whitney test), ATP (300 μM) (*P* = 0.770, unpaired *t* test) and NE (20 μM) (*P* = 0.730, unpaired *t* test). (**K**) Quantification of relative mRNA levels of mGluR3 (*P* = 7.2E-10, unpaired *t* test) and mGluR5 (*P* = 3.4E-7, unpaired *t* test) by RT-qPCR analyzing of GLAST^+^ MACS cells (n = 3 independent preparations). \**P* < 0.05, \*\**P* < 0.01, \*\*\**P* < 0.001, n.s, not significant. Data are shown as mean ± s.e.m.

The assessment of acute effects of Sevo on astrocyte Ca^2+^ signals was achieved by continuous infusion of Sevo (0.36 ± 0.03 mM, equivalents to 2.5% Sevo (18) in acute brain slices from WT mice at P14. Interestingly, we found that acute Sevo exposure led to a gradual and persistent loss of basal Ca^2+^ level in cortical developing astrocytes, and the effect was not driven by action potential firing, as basal Ca^2+^ level still decreased in the presence of 300 nM tetrodotoxin (TTX) (**Fig. 3B, C**). Independently, we performed GCaMP6f microinjections at P0, Sevo/Control exposure *in vivo* at P7, and we found that the GCaMP6f basal fluorescent intensity in Sevo group mice was significantly lower than the Control group mice (**Fig. 3D, E**) when measured at P14, suggesting the loss of basal Ca^2+^ level was not fully recovered within 6-8 days after exposure. To explore if the long-term loss of basal Ca^2+^ level is age-dependent, we performed similar experiments in adult animals. Acute Sevo exposure also led to a similar extent of decrease in basal Ca^2+^ level in mature astrocytes (**Supplementary Fig. 7A**) compared the developing ones. However, when performed Sevo/Control exposure *in vivo* at P56, we found no significant difference in GCaMP6f basal fluorescent intensity in the two group mice measured at 7 days later at P63 (**Fig. 3E**). Together, these results suggest mature astrocytes are more resistant in response to the decrease of basal Ca^2+^ level than the immature ones following lengthy general anesthesia.

Spontaneous Ca^2+^ signals were analyzed automatically with post manual inspection by using the computational tool suite “Functional AStrocyte Phenotyping (FASP)” (19). Similar to previous studies in mature cortical astrocytes (20), developing cortical astrocytes displayed waves (events area >10 μm^2^) and microdomain signals (events area 1.5-10 μm^2^). Consistent with the previous work (17), acute Sevo exposure resulted in decreased frequency and amplitude of Ca^2+^ waves (**Supplementary Fig. 7B**). In addition, Sevo exposure *in vivo* at P7 also resulted in altered spontaneous Ca^2+^ signals properties measured at P14, including amplitude, half width (**Fig. 3F-H**). We also assessed neurotransmitter-evoked Ca^2+^ signals and found the peak amplitude of mGluR2/3 agonist LY354740-evoked Ca^2+^ signals and mGluR1/5 agonist DHPG-evoked Ca^2+^ signals were significantly lower in Sevo group mice. While ATP-evoked and Noradrenaline (NE)-evoked Ca^2+^ signals were similar between these two groups (**Fig. 3I, J**). We further evaluated the expression of mGluR3 and mGluR5 in cortical astrocytes at P14, by using RT-qPCR for magnetic cell sorting (MACS) isolated cortical astrocytes (GLAST^+^ cells) from Control and Sevo group mice (**see methods**). We found that both mGluR3 and mGluR5 mRNA levels were significantly lower in Sevo group mice (**Fig. 3K**).

Together, this dataset suggests that lengthy Sevo exposure disrupted both acutely and chronically the basal Ca^2+^ levels, spontaneous and neurotransmitter-evoked Ca^2+^ transients in developing cortical astrocytes.

### Down-regulation of Ezrin expression with lengthy Sevo exposure

Ezrin is an actin-binding membrane-bound protein expressed mainly within astrocyte processes in the central nervous system (21), and was found required for the structural plasticity of astrocyte processes in culture (22). Since the loss of morphology in Sevo group mice predominantly occurred in the fine processes (**Fig. 1**), we hypothesized that Ezrin may be a key structural determinant of astrocyte fine processes *in vivo*, and its dysfunction may occur following lengthy general anesthesia. To this end, first, we found that, in the somatosensory cortex of P21 mice, Ezrin was only partially colocalized with the astrocyte marker S100β, which labeled mainly the soma and primary branches (**Fig. 4A**), whereas most of the Ezrin signals seemed to be peripheral to S100β signals. To precisely quantify the distribution of Ezrin within astrocyte territory, we fluorescently labeled cortical astrocytes by stereotactic injection of AAV5⋅gfaABC1D⋅mCherry into the cortex of P0 mice and stained Ezrin at P9, P16 and P21 (**Fig. 4B, C**). We found that the volume fraction of Ezrin was gradually increased in fine processes but decreased in soma and primary branches of cortical astrocytes from P9 to P21 **(Fig. 4C, D)**. At P21, more than 70% of total Ezrin was localized within astrocyte fine processes. Interestingly, the expression of Ezrin in the somatosensory cortex, but not in hippocampal CA1sr or DG, was significantly down-regulated in Sevo group mice compared to Control group mice at P14 (**Fig. 4E, F; Supplementary Fig.8**). Consistent with immunostaining data, mRNA level of Ezrin in the somatosensory cortex of Sevo group mice at P8 and P14 were also significantly lower, detected by using RT-qPCR for GLAST^+^ MCAS cells (**Fig. 4G**). Together our data suggest that Ezrin was developmentally enriched within astrocyte fine processes, and was down-regulated in Sevo group mice.

**Fig 4.**
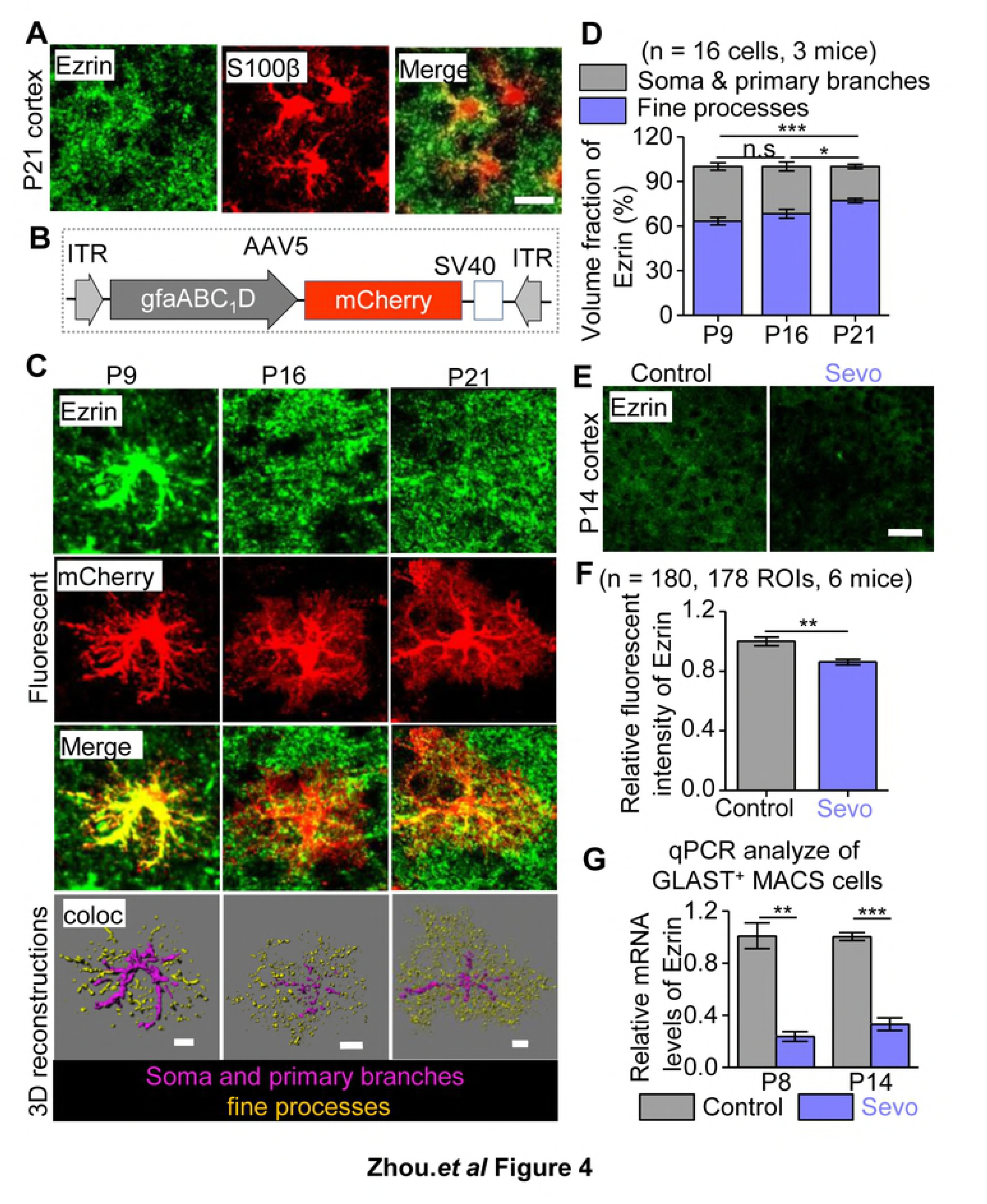
Down-regulation of Ezrin expression with lengthy Sevo exposure. (**A**) Representative images of Ezrin colocalized with S100β. Scale bars, 10 μm. (**B**) Schematic of construction of AAVs. (**C**) Representative fluorescent images of Ezrin and mCherry labeled astrocytes, and 3D reconstruction of Ezrin within astrocytic fine processes or soma and primary branches in the somatosensory cortex at P9, P16 and P21. Scale bars, 10 μm. (**D**) Volume fraction of Ezrin within the fine processes, soma and primary branches of astrocytes (P9 *vs* P16: *P* = 0.304, P16 *vs* P21: *P* = 0.04, P9 *vs* P21: *P* = 6.5E-4, one-way ANOVA test; *post-hoc* Tukey test). (**E**) Confocal images for Ezrin staining. Scale bars, 50 μm. (**F**) Quantification of Ezrin fluorescent intensity (*P* = 0.007, Mann-Whitney test). (**G**) Quantification of Ezrin mRNA levels of GLAST^+^ MACS cells in the cortex of Control and Sevo group mice at P8 (n = 3 independent preparations, *P* = 0.002, unpaired *t* test) and P14 (n = 3 independent preparations, *P* = 0.002, unpaired *t* test). \*\**P* < 0.01, \*\*\**P* < 0.001, n.s, not significant. Data are shown as mean ± s.e.m.

### Astrocyte morphological deficits were induced by down-regulation of Ezrin in a Ca^2+^ dependent manner

We then asked whether the down-regulation of Ezrin by early Sevo exposure was due to the disrupted astrocyte Ca^2+^ signals (**Fig. 3**). Intracellular Ca^2+^ chelation was achieved by incubating cultured astrocytes with a membrane-permeant Ca^2+^ chelator BAPTA-AM (30 μM). The Ezrin expression in the primary cultured astrocytes was significantly decreased after 1 h-incubation with BAPTA-AM **(Fig. 5A, B)**. Interestingly, BAPTA-AM (30 μM, 30 min) incubation induced marked morphological changes, with decreased surface area and perimeter, revealed by successive imaging of enhanced green fluorescent protein (eGFP) plasmid transfected astrocytes *in vitro* (**Fig. 5C-E**). Similar morphological deficits were also observed in astrocytes in cortical slices treated with BAPTA-AM, where AAV5⋅gfaABC_1_D⋅eGFP was microinjected into the mouse cortex at P0 to sparsely label astrocytes *in vivo* (**Fig. 5F**). Astrocytes fine process volume was decreased after BAPTA-AM incubation for 1 h in the presence of TTX (**Fig. 5G, H**). Thus, our data so far suggest that reducing astrocyte intracellular Ca^2+^ was sufficient to produce the loss of Ezrin and the loss of astrocyte morphology.

**Fig 5.**
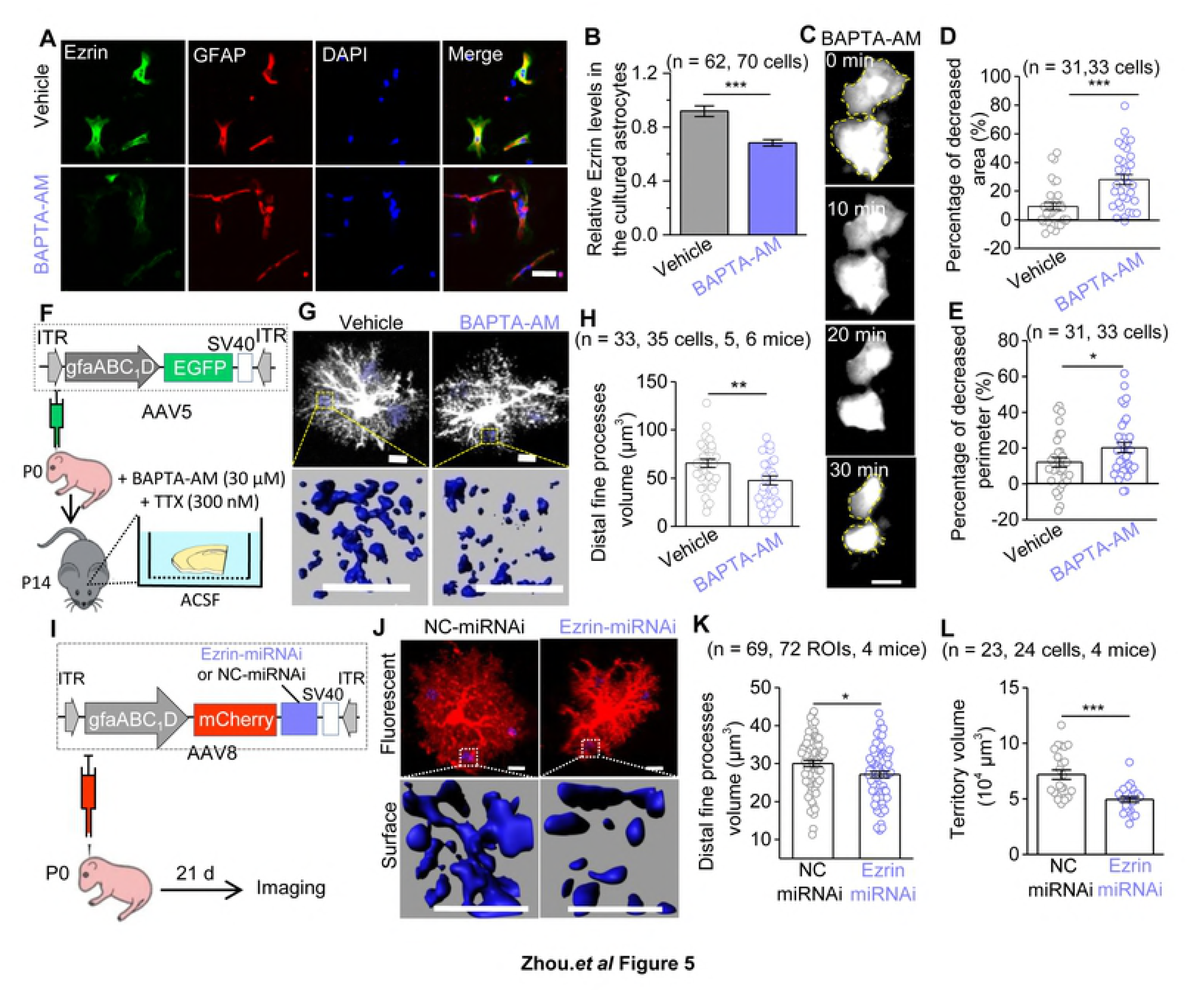
Reducing astrocyte intracellular Ca^2+^ leads to the down-regulation of Ezrin, which produces morphological deficits. (**A**) Representative immunofluorescence images of Ezrin in the primary cultured astrocytes after 1 h-incubation with Vehicle (DMSO) or BAPTA-AM (30 μM). Scale bars, 50 μm. (**B**) Quantification of relative fluorescent intensity of Ezrin (n = 3 independent preparations; *P* = 4.3E-6, Mann-Whitney test). (**C**) Representative successive imaging of EGFP labeled astrocytes during BAPTA-AM (30 μM) incubation. Scale bars, 50 μm. (**D, E**) Quantification cultured astrocytes morphology (n = 3 independent preparations; **D**: *P* = 8.2E-5, Mann-Whitney test; **E**: *P* = 0.041, unpaired *t* test). (**F**) Diagram illustrates the vector construction, AAV injection in neonate mice and protocol for brain slices incubation. (**G**) Representative fluorescent images and 3D reconstructed distal fine processes of astrocytes within a 10 × 10 × 10 μm^3^ ROI, 3 ROIs for one cell. Scale bars, 10 μm. (**H**) Quantification of astrocyte distal fine processes (*P* = 0.006, unpaired *t* test). (**I**) The vector construction of AAV8⋅gfaABC_1_D⋅mCherry⋅Ezrin.miRNAi (or NC-miRNAi) and stereotactic injection of AAVs into P0 mice. (**J**) Confocal images and 3D reconstructed astrocytes. (**K**) Quantification of astrocytic distal fine processes volume in NC-miRNAi and Ezrin-miRNAi mice at P21 (*P* = 0.018, unpaired *t* test). Scale bars, 10 μm. (**L**) Quantification of astrocyte territory volume (*P* = 5.0E-5, unpaired *t* test). \**P* < 0.05, \*\**P* < 0.01, \*\*\**P* < 0.001. Data are shown as mean ± s.e.m.

Did the loss of Ezrin lead to the loss of astrocyte morphology? We sought to genetically knock down Ezrin within cortical astrocytes *in vivo* using microRNA silencing delivered by AAVs. We designed AAV8⋅gfaABC1D⋅mCherry⋅Ezrin-miRNAi (Ezrin-miRNAi) and its negative control AAV8⋅gfaABC1D⋅mCherry⋅NC-miRNAi (NC-miRNAi) (**see methods**), and performed microinjections into the cortex of P0 mice (**Fig. 5I**). Ezrin-miRNAi injection resulted in a significantly down-regulation of Ezrin expression (by ~26%) in the somatosensory cortex of mice at P21, assessed by Ezrin immunostaining (**Supplementary Fig. 9A, B**). The miRNAi expression is highly specific to astrocytes, as 94.4% of mCherry^+^ cells colocalized with S100β and only 5.6% of which colocalized with NeuN **(Supplementary Fig. 9C, D)**. Furthermore, Ezrin-miRNAi or NC-miRNAi injection into the cortex of P0 mice did not cause apparent astrogliosis at P21 (**Supplementary Fig. 9E**). As expected, Ezrin knocking-down resulted in decreased astrocytes fine processes and territory volume in P21 mice (**Fig. 5J-L**), whereas the soma volume was slightly increased and primary branches volume and number were unchanged (**Supplementary Fig. 9F, G**).

Together, our data suggest that suppressing intracellular Ca^2+^ signals in astrocytes produced severe astrocyte morphological deficits along with the down-regulation of Ezrin, and importantly, knocking-down Ezrin *in vivo* was sufficient to produce the loss of astrocyte fine processes, which was very similar to the structural loss of the cortical astrocytes from Sevo group mice.

### Down-regulation of Ezrin resulted in increased synapses with decreased synaptic functions

In the last set of experiments, we tested the hypothesis that the compromised astrocyte morphogenesis mediated by the down-regulation of Ezrin drove the aberrant excitatory synaptic structure and function in Sevo group mice. In mice with Ezrin-miRNAi or NC-miRNAi microinjection at P0, we quantified the synaptic density within the territories of Ezrin-or NC-miRNAi expressing astrocytes (which are mCherry^+^) and compared them with those are outside (which are mCherry^−^) around P21. Synapses were identified by the co-localization of presynaptic marker VGluT1 and postsynaptic marker PSD95. We found that the density of excitatory synapses within the mCherry^+^ area was significantly higher when compared with the mCherry^−^ area in Ezrin-miRNAi mice, whereas in NC-miRNAi mice, no significant difference was found between the mCherry^+^ and the mCherry^−^ area (**Fig. 6A, B**), suggesting Ezrin knock-down resulted in an increase of the excitatory synapses within the astrocyte territory. We next recorded the mEPSCs and eEPSCs in L 3-5 pyramidal neurons from Ezrin-miRNAi and NC-miRNAi mice. The mEPSCs frequency was significantly 50% smaller in Ezrin-miRNAi mice compared to NC-miRNAi mice (**Fig. 6C, D**), while the mEPSCs amplitude was similar between the two groups (**Fig. 6E**). The AMPAR/NMDAR ratio was significantly 35% smaller in Ezrin-miRNAi mice (**Fig. 6F, G**). The similar extent of reductions in eEPSCs frequency and AMPAR/NMDAR ratio suggest the decreased synaptic transmission are largely AMPA-dependent. Together, the structural and functional data suggest that astrocytic Ezrin knock-down was sufficient to produce an increase of the synaptic density, with reduced AMPA-mediated synaptic transmission, a phenotype virtually identical to the one found in the cortex of Sevo group mice (**Fig. 2**).

**Fig 6.**
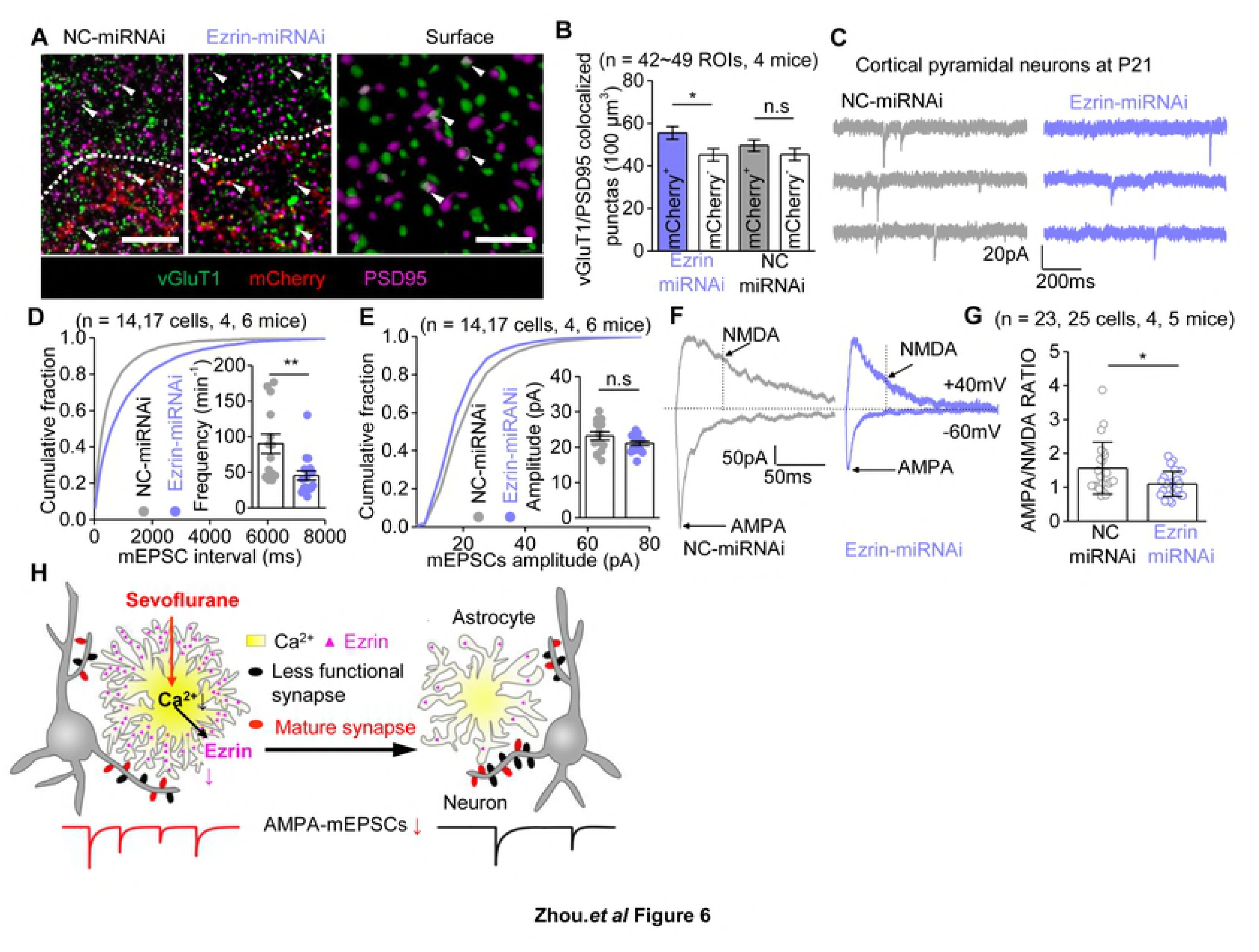
Down-regulation of Ezrin drives increased total synapses with decreased synaptic function. (**A**) Representative confocal images and 3D reconstruction of vGluT1 (green) and PSD95 (magenta). The mCherry^+^ (red) and mCherry^−^ regions are separated by dash line and vGluT1/PSD95 colocalized punctas are shown (arrow). Scale bars, 5 μm (left), 1 μm (right). (**B**) Quantification of vGluT1/PSD95 colocalized puncta density (per 100 μm^3^) in the mCherry^+^ and mCherry^−^ regions of Ezrin-miRNAi (*P* = 0.014, Mann-Whitney test) and NC-miRNAi (*P* = 0.288, unpaired *t* test) injected mice at P21. (**C**) Representative mEPSCs traces. (**D**) Cumulative distributions and average mEPSCs frequency (*P* = 0.005, Mann-Whitney test). (**E**) Cumulative distributions and average of mEPSCs amplitude (*P* = 0.108, unpaired *t* test). (**F**) Representative eEPSCs traces. AMPAR-mediated synaptic currents were recorded with the presence of 100 μM picrotoxinin and 20 μM glycine, and mixed AMPA and NMDA currents were recorded with 100 μM picrotoxinin and 20 μM glycine. The black dotted line (50 ms after the stimulation) indicates measuring points for NMDAR-eEPSC amplitudes. (**G**) Quantification of AMPA/NMDA ratio. (*P* = 0.018, Mann-Whitney test). **(H)** Summary for the key findings in the current study. \**P* < 0.05, \*\**P* < 0.01, \*\*\**P* < 0.001, n.s, not significant. Data are shown as mean ± s.e.m.

## Discussion

There are several key findings from this study, some of which are schematized in **Fig. 6H.** First, we reported in details how lengthy Sevo exposure disrupts astrocyte morphogenesis. Compromised astrocyte morphogenesis was associated only with a relatively long exposure (4 h), and occurred only in the developing brain but not in the mature brain. Second, we showed that the Sevo-induced aberrant synaptogenesis and astrocytic morphological deficits were spatiotemporally correlated. Third, we revealed the underlying mechanisms for compromised astrocyte morphogenesis, in which Sevo targeted astrocyte Ca^2+^ signaling, and hereby disrupted Ezrin expression, which functioned as a key regulator of astrocyte morphogenesis *in vivo*. We further showed Ezrin was developmentally up-regulated in astrocyte fine processes, and the disruption of which predominantly resulted in the loss of the fine processes. Fourth, we showed that, in WT mice, the down-regulation of Ezrin within astrocytes was sufficient to produce an increase of synapses accompanied with a decrease of AMPA-mediated synaptic functions, a phenotype virtually identical to the one found in the Sevo group mice, which strongly suggests that Ezrin-mediated compromised astrocyte morphogenesis likely drove the dysfunctional synaptogenesis in the developing cortex with lengthy Sevo anesthesia.

Apoptosis and abnormal neural circuit formation including aberrant neurite growth, guidance and synaptic connections are the major molecular and cellular phenotypes reported in the GAs-induced neurotoxicity (2). In the current study, we focused on the structural and functional deficits in astrocytes and neurons that are highly relevant to the neural circuit formation and functions. The initial goal was to evaluate how lengthy Sevo exposure induces astrogliosis. To our surprise, in the Sevo group mice, cortical astrocytes displayed loss of fine processes without apparent astrogliosis. The loss or the delayed growth of astrocyte fine processes, although subtle, was identified using two independent methods: single astrocyte fluorescent labeling and SBF-SEM. At P21, structural loss in astrocytes was not visible under light microscopy, but remained profound under SBF-SEM. Our study indicates that the detailed morphological analysis of astrocytes in addition to conventional staining for astrogliosis or cytoskeletal impairment is needed for a better understanding of astrocyte biology in the context of diseases or injuries.

Astrocyte morphological deficits displayed exposure-length-, age- and brain region-dependence. The exposure-length and age dependence are in consistent with the clinical findings so far, where only infants or children with repeated or prolonged general anesthesia are associated with cognitive dysfunctions. Consistent with a previous animal study with propofol (23), we found Sevo lengthy anesthesia induced an increase of dendritic spines and synapses at P21 in the cortex, but decreases were also reported (24, 25). We did not observe significant astrocyte nor neuron structural deficits in developing hippocampal CA1sr and DG-mo regions, whereas a few studies found abnormal synaptic growth (26), altered synaptic plasticity (24) in the hippocampus. The inconsistency may due to different anesthetic agents (isoflurane, sevoflurane, propofol etc.) used, different exposure conditions (different concentration and exposure time, repeated exposure or single exposure) or different type of cells examined (pyramidal neurons or granule cells) or any other different experimental settings. Regardless, with experiments performed with intact tissue preparation or *in vivo* under our experimental paradigms, we revealed both neuronal and astrocytic deficits that were highly correlated in the somatosensory cortex, and likely in other cortical areas.

Astrocytes use intracellular Ca^2+^ signals as one of their primary forms to engage in neural circuits, and the disruption of which alters behavior (27) and contributes to diseases and injuries (28). We found that Sevo disrupted astrocyte intracellular Ca^2+^ transients and basal Ca^2+^ level both acutely and chronically. Interestingly, most of the clinically used GAs suppress Ca^2+^ transients *in vivo* (17), therefore, it is likely that astrocytes Ca^2+^ may represent a common pathway for GAs-induced neurotoxicity. Further experiments are needed to test if other GAs with distinct action mechanisms, including isoflurane, propofol and ketamine, disrupt Ca^2+^ homeostasis in developing astrocytes. The decrease of basal Ca^2+^ level by Sevo is of particular interest, since it has been recently shown that basal Ca^2+^level decreased from P15 to P21 in mouse hippocampus, suggesting there is an intrinsic regulatory mechanism for basal Ca^2+^ level during development (29). The molecular pathway(s) by which Sevo disrupts astrocyte Ca^2+^ efflux/influx is currently unknown. A wide range of ion channels and transporters have known sensitivities to inhaled anesthetics (30), some of which are expressed in astrocytes, such as sodium–calcium exchangers (NCX), nicotinic receptors, background two-pore domain K^+^ channels etc. The net effects of Sevo on astrocyte Ca^2+^ likely involve multiple targets and multiple mechanisms, which require further studies to clarify.

Actin-binding proteins of the Ezrin, Radixin, and Moesin (ERM) family play key roles in the molecular mechanisms of cell motility, process formation, and tumorigenesis. In central nervous system, Ezrin was found specifically localized to fine astrocytic processes (21, 22). We extended the previous findings by showing that the development profile of Ezrin expression was correlated with astrocyte morphological maturation, a process involves in a substantial growth of the fine processes, and we provided direct evidence for the functional involvement of Ezrin in astrocyte fine structure growth by showing that the disruption of Ezrin *in vivo* predominantly affected the formation of the fine processes. Astrocyte fine processes display actin-dependent structural plasticity that is regulated by synaptic activity through astrocyte metabotropic glutamate receptors and intracellular Ca^2+^ signaling, and the astrocyte structural plasticity in turn controls synapse stability (31). Therefore, in the subsequent studies, it would be of particular interest to test if Ezrin is involved in neural-activity dependent structural plasticity at the synaptic scale, thereby participates in astrocyte-neuron dynamic interactions *in vivo*, and ultimately, in behavior.

Both lengthy Sevo exposure and Ezrin knock-down within astrocytes produced an intriguing and apparently paradoxical neuronal phenotype: an increase of total synapses with reduced AMPA-mediated synaptic function. The most straightforward explanation is that there was a much greater proportion of silent synapses in Sevo and Ezrin-miRNA group mice, whereas the proportion of functional ones was reduced. However, we could not exclude a decrease of presynaptic glutamate release probabilities in both Sevo and Ezrin-miRNA group mice. Here we focus our discussion on astrocytic roles in forming functional and nonfunctional synapses. In the developing brain, newly formed glutamatergic synapses are often associated with silent post-synaptic spines that lack functional AMPA receptor-mediated transmission. Most of these silent synapses are eliminated, but some are specifically selected for AMPA un-silencing and lead to stabilization during the development (32). Increased (less-functional) synapses with reduced functions likely resulted from an imbalance among all the three key processes required for proper synaptogenesis: synaptic formation, stabilization, and elimination. There are a few astrocyte-based mechanisms for synaptogenesis. For instance, astrocyte-secreted molecules including thrombospondins and hevin promote structurally normal but functional silent synapses, while glypican 4 and glypican 6 convert silent synapses into functional ones by increasing the surface level of AMPAR, and MEGF10 and MERTK mediate synapse elimination through phagocytosis (33). All the signaling pathways above require proper spatial interactions between neurons and astrocytes. We hypothesize that astrocyte structural deficits resulted in an increase or decrease of astrocyte–neuron signaling efficiency due to the altered spatial interactions between astrocytes and neurons at the synaptic level. In support, recent work by Stogsdill *et al* (34) demonstrated that astrocyte–neuron adhesions control astrocyte morphogenesis and thereby control synaptogenesis.

Unexpectedly, our data reveal that astrocytes play a central role between GAs and neurotoxicity in the developing brain. Postoperative cognitive dysfunction (POCD) is a chronic decline in cognitive function also associated with anesthesia/surgery. It was found that POCD was associated with astrocytic decreased Ca^2+^ signaling along with enhanced synaptic transmission in the mouse hippocampus (35). The causal relationship between astrocyte and neuron dysfunctions in POCD remains to be established. More broadly, in addition to the earlier studies in Huntington’s disease (HD) mouse models where astrocytes displayed functional (36) and morphological (15) deficits before astrogliosis, our study presented here reinforces the idea that astrocyte fine structural integrity may represent a therapeutic target in a variety of neurological and psychiatric diseases.

## Materials and Methods

### Ethics statement

All experiments with mice were approved by the Animal Research Committee at the West China Hospital of Sichuan University (protocol 2018159A). For tissue collection, mice were given a lethal dose of pentobarbital intraperitoneally.

### Animals

C57BL/6 mice were housed in a temperature- and humidity-controlled room with a 12-h light–dark cycle, and provided with *ad libitum* access to water and food. Both sexes were equally presented in all experiments.

### Sevoflurane exposure protocol

P7 mouse littermates were randomly assigned to 2 groups. In Sevo group, mice were exposed to 2.5% sevoflurane (HENGRUI MEDICINE) carried in 30% O_2_/70% CO_2_ at a flow rate of 1.5 L/min for 4 h. An agent-specific vaporizer (Vanbon, ibis 200) was used to deliver sevoflurane. Gas was monitored by an agent gas monitor (Philips). Mice were continually monitored and recorded for skin temperature and respiratory rate during anesthesia. Animals were returned to their mothers upon regaining righting reflex. Intracardiac puncture was used to collect left ventricular blood samples for arterial blood gas analysis (Mindray, BC-3000).

### Intracellular Lucifer yellow dye filling and morphological 3D reconstructions

#### Astrocytes

Intracellular dye filling was performed on the lightly fixed brain slices from the layer 3-5 (L3-5) of primary somatosensory cortex, hippocampal CA1 sr and hippocampal DG of mice at P8, P14 and P21 using the protocols as previously reported. After injection, the confocal imaging stacks were collected with a Z-step size of 0.25 µm under a multiphoton microscope (Nikon A1R^+^). 3D reconstructions were processed offline using Imaris 7.4.2 (Bitplane, South Windsor, CT, USA) as reported previously (15). To quantify the fine processes of astrocytes, six randomly chosen 5 µm × 5 µm × 5 µm ROIs devoid of the soma and large branches were reconstructed with the *Local Contrast* method. Three of these ROIs that near the soma were termed as proximal fine processes, and the other three ROIs that located in the edge of astrocyte were termed as distal fine processes. 3D surface rendering of astrocytes for calculating the territory volume was achieved similarly, utilizing *Background Subtraction* of Imaris.

#### Dendritic spines

Dendritic spines visualization using intracellular Lucifer yellow dye filling was performed similarly as previously described (37). Consecutive stacks of images with Z-step of 0.25 μm were acquired at high magnification (60× glycerol, 5× optical zoom) using a multiphoton microscope (Nikon A1R^+^). The image stacks were then reconstructed and analyzed by using the *Filament function* of Imaris. *Dendritic spine density* was determined by counting the number of dendritic spines per 10 μm of dendritic length, starting from 50 μm to 120 μm away from the soma, where the dendritic spine density was reported with low variety. For apical dendrites, the 70 μm long segment was also analyzed every 10 μm of dendritic length. For spine classification, the *Filament function* of Imaris was used; stubby: length (spine) < 1.5 and max width (head) < mean width (neck) ×1.2; mushroom: max width (head) > mean width (neck) × 1.2 and max width (head) > 0.3; the rest were defined as long-thin (26). We only quantified the total and mushroom spine density, but the stubby and long thin were difficult to be classified in many cases due to the limited image resolution.

### Serial block face scanning electron microscopy

Mice were euthanized with 100 mg/kg pentobarbital and transcardially perfused with 40 ml of fixative solution (2% paraformaldehyde Sigma-Aldrich #158127) and 2.5% glutaraldehyde (Sigma-Aldrich #G7651) in a 0.1 M Phosphate (PB) buffer, pH 7.4. Brains were sliced into 100 μm coronal sections and we then further dissected the L3-5 of primary somatosensory cortex. Tissues were post-fixed in fixative solution for 48 h post euthanasia at 4°C. After washing by PB (pH 7.4) for three times, tissues were fixed with osmium-ferricyanide, followed by thiocarbohydrazide treatment, and then further fixed with 1% aqueous osmium tetroxide. Samples were then incubated overnight in 2% aqueous uranyl acetate at 4°C. Next, samples were dehydrated through a series of alcohol solutions and embedded in SPI-Pon 812-substitute resin. The block was carefully trimmed for focused ion beam scanning electron microscopy imaging with Helios NanoLab 600i (FEI). Samples were imaged at accelerated voltage 4 keV, beam current 0.34 nA, at 3.37 nm/pixel resolution, with horizontal field width 13.8 μm and slice thickness of 60 nm (38). Each image series contains 90-120 sections with a total volume of 3313.3-3542.2 μm^3^ in Sevo and Control group at P14 and P21. Image series were registered and then analyzed using *Reconstruct* software version 1.1 (https://synapseweb.clm.utexas.edu/software-0). Astrocytic profiles are clearly identifiable in serial images owing to their pale cytoplasm, irregular contouring border and the absence of vesicles as well as synapses. Only asymmetric synapses identified by the shape of their axonal and dendritic spines and their synaptic vesicles were included in the quantification. We randomly selected fields of 16 μm^2^ and thickness of 1.8 μm (volume 28.8 μm^3^) or fields of 25.1 μm^2^ and thickness of 3 μm (volume 75.3 μm^3^) from blocks in Sevo and Control group mice for reconstruction. These volumes did not include any large dendritic profiles or soma of neurons, glia or endothelial cells. To quantify astrocyte– synaptic cleft contacts, only astrocytic process endings visually directly apposed postsynaptic densities (PSD), i.e. with no intervening elements of the neuropil, were counted. Cross-sectioned astrocyte perimeters of the synaptic cleft were computed by assigning the contact point between the astrocytic membrane and the synaptic cleft in individual sections then summing over all sections. The mushroom dendritic spines were recognized as the presence of a spine apparatus and a complex PSD (perforated, U-shaped or segmented) whereas other spines had only macular PSDs and no spine apparatus (39). The stubby spines were not separately classified from long thin spines.

### AAVs generation

The pZac2.1.gfaABC_1_D⋅cyto-GCaMP6f plasmid was a gift form Dr. Baljit Khakh (UCLA), and was send to Dr. Biao Dong’s Lab (Sichuan University) for AAV package. The AAV5⋅gfaABC_1_D⋅eGFP and AAV5⋅gfaABC_1_D⋅mCherry were gifts from Taitool Bioscience. To achieve astrocyte specific Ezrin knock-down using microRNA based silencing technique, we used the BLOCK-iT™ Pol II miR RNAi Expression Vector Kits (Invitrogen). Six pre-miRNA sequences for Ezrin (Ezrin-miRNA) and negative control sequence (NC-miRNA) were designed (Invitrogen’s RNAi Designer), synthesized and cloned into pAAV⋅CMV_bGI⋅mCherry⋅miRNAi vector (Taitool Bioscience). The knockdown (KD) efficiency were then evaluated by cotransfecting eGFP-tagged-Ezrin with the Ezrin miRNA vectors separately in human embryonic kidney (HEK293) cells and the KD efficiency were indicated by reduction of fluorescence signal expressed by eGFP-Ezrin vector. The most effective sequence was chosen as follow: Ezrin-miRNA: 5’-AAAGTCAGGTGCCTTCTTGTC-3’, and NC-miRNA: 5’-AAATGTACTGCGCGTGGAGAC-3’ The selected oligos were then cloned into the linearized pAAV⋅gfaABC_1_D⋅mCherry⋅miRNAi vector (Taitool Biosciense) using T4 DNA ligase. The plasmids were packaged into AAV8 virus by calcium phosphate transfection with capsid and helper vectors on HEK293 cells. The collected viruses were purified by iodixanol density gradient centrifugation. The titer of AAV8⋅gfaABC_1_D⋅mCherry⋅Ezrin-miRNAi and AAV8⋅gfaABC_1_D⋅mCherry⋅NC-miRNAi was determined by qPCR.

### AAVs microinjections

P0 neonates were under cryo-anesthesia before injection as previously reported (40). Viral injections were performed by using a stereotaxic apparatus (RWD) to guide the placement of Hamilton syringe fixed with beveled glass pipettes (Sutter Instrument, 1.0 mm outer diameter) into the cortex. The injection site was located a half of the distance along a line defined between each eye and the lambda intersection of the skull. The needle was held perpendicular to the skull surface during insertion to a depth of approximately 0.2 mm. 1.0 µl of AAV5⋅gfaABC_1_D⋅cyto-GCaMP6f (GCaMP6f) (2.0 × 10^13^ gc/ml) or 1.0 µl of AAV5⋅gfaABC_1_D⋅mCherry (3.46 × 10^12^ gc/ml) or 1.0 µl of AAV5⋅gfaABC_1_D⋅EGFP (2 × 10^13^ gc/ml) or 0.6 µl of Ezrin-miRNAi (5.5 × 10^12^ gc/ml) or 0.6 µl of NC-miRNAi (5.4 × 10^12^ gc/ml) was slowly injected into both sides of the hemisphere. Glass pipettes were left in place for at least 5 min. After injection, pups were allowed to completely recover on a warming blanket and then returned to the home cage.

### Acute brain slice preparation and electrophysiology

Acute slices were prepared as previously described (41). L3-5 pyramidal neurons in somatosensory cortex in slices were visualized with infrared optics on an upright microscope (BX51WI, Olympus). Clampex 10.4 software and a MultiClamp 700B amplifier (Molecular Devices) were used for electrophysiology (Molecular Devices). All recordings were carried out at room temperature. Recordings were filtered at 2 KHz, digitized at 10 KHz and acquired with Digidata 1440A (Molecular Devices). The recording pipettes (3-4 MΩ) were pulled from borosilicate glass by using a Flaming-Brown horizontal puller (Model P97, Sutter Instruments). For mEPSCs recordings, the intracellular solution in the patch pipette contained the following (in mM): 130 KCl, 2 NaCl, 10 HEPES, 5 EGTA, 2 Mg-ATP, 0.5 CaCl_2_, pH 7.3 adjusted with KOH. For eEPSCs recordings, the intracellular solution in the patch pipette contained the following (in mM): 120 Cesium methanesulfonate, 15 CsCl, 8 NaCl, 10 HEPES, 0.2 EGTA, 10 TEA-Cl, 2 Mg-ATP, 0.3 Na_2_-GTP, 3 QX-314, pH 7.3 adjusted with CsOH. The initial access resistances were < 20 MΩ for all cells; if this changed by > 25%, the cell was discarded. Synaptic currents were collected for 3-5 min for each cell. Analysis was carried out by using Mini Analysis Program (version 6.0.3; Synaptosoft, Leonia, NJ). Only events greater than 8 pA, with a rise time less than 2 ms and decay time less than 10 ms were included in the analysis. For eEPSCs, a bipolar stimulating electrode was placed in the somatosensory cortex. The recording pipettes were typically located 250–300 μm away from the stimulation site. Only single-peaked responses were included for analysis. The time constant for the decay of eEPSCs was determined by fitting the decay to a single exponential by using pCLAMP10.7 software.

### Ca^2+^ imaging

Acute brain slices from mice injected with GCaMP6f were prepared for Ca^2+^ imaging. Slices were imaged using a Nikon A1R^+^ multiphoton microscope with 40x (NA 0.8) water immersion objective (Nikon), using the 488 nm laser. Slices were continuously superfused at 1-2 mL/min with oxygenated aCSF at room temperature. Images were typically 512 × 512 pixels/frame with 1-2× optical zoom at a scan rate of 1 sec per frame. Drugs (LY354740:10 μM, Tocris; DHPG; 100 μM, Tocris; ATP: 300 nM, Sigma-Aldrich; NE: 20 μM, Abisin) were loaded by bath application. For acute administration of sevoflurane, 2.5% sevoflurane was equilibrated in bathing solutions in a reservoir by passing air (flow rate, 0.5 L/min) through a calibrated vaporizer for at least 30 min before entering a recording chamber. Samples of the superfusate in the recording chamber were collected for the measurement of the sevoflurane concentration by gas chromatography. The mean sevoflurane concentration was 0.36 ± 0.03 mM. Recordings were excluded from analysis if the cell drifted out of frame or out of the z-plane. Lateral drifts in astrocyte position were corrected with the TurboReg Plugin in ImageJ. Spontaneous Ca^2+^ signals were analyzed by MATLAB (R2015a, MathWorks) using custom-written scripts called “Functional Astrocyte Phenotyping (FASP)” (19). A signal was declared as a Ca^2+^ transient if it exceeded the detection by greater than twice of the baseline noise (SD).

### Tissue dissociation, astrocyte sorting and qPCR

The cortical hemispheres from Control and Sevo group mice at P14 were dissociated following published guidelines (42) with slight modifications. Briefly, the cortex from 4 mice were dissected and digested together for 45 min at 37°C with 10 ml of enzyme solution (Dulbecco’s minimum essential medium (DMEM), 50 mM EDTA, 50 U/ml DNase-1, 0.1% collagenase and 0.05% trypsin) while bubbling with 5% CO_2_/95% O_2_. After digestion, the tissue was mechanically dissociated and filtered with a 70 μm mesh. Astrocytes were separated by a magnetic cell sorting (MACS) method according to the manufacturer’s protocol (Miltenyi Biotec, The Netherlands). Cell suspensions were labeled with superparamagnetic MicroBeads coupled to antibodies specific for the astrocyte marker GLAST (Anti-GLAST, MicroBead Kit, Miltenyi Biotec). Before antibody labeling, nonspecific binding to the Fc receptor was blocked using the FcR Blocking Reagent (Miltenyi Biotec). Cells were suspended in PBS with 0.5% BSA and the cell suspension was loaded onto an MS Column (Miltenyi Biotec), which was placed in the magnetic field of a MiniMACS^TM^ Separator (Miltenyi Biotec). The magnetically labeled GLAST-positive cells were retained within the column and eluted as the positively selected cell fraction after removing the column from the magnet. RNA from the sorted cells was extracted and converted to cDNA. qPCR was performed as reported previously (41).

Sequences of primers used for RT-PCR or qPCR (5’ to 3’)

*Grm3* (PrimerBank ID, 32469489a1): For-CTGGAGGCCATGTTGTTTGC

Rev-CATCCACTTTAGTCAACGATGCT

*Grm5* (PrimerBank ID, 219801746c1): For-ACCAACCAACTGTGGACAAAG

Rev-CAAGAGTGTGGGATCTGAATTGA

*Ezr* (PrimerBank ID, 83921617c2): For-CACAGGAGGTCCGAAAGGAGA

Rev-CTTGGCCTGAACGGCATAGG

*gapdh* (PrimerBank ID, 126012538c1): For-AGGTCGGTGTGAACGGATTTG

Rev-GGGGTCGTTGATGGCAACA

### Primary astrocyte culture

Primary astrocyte cultures were prepared from the cerebral cortex of newborn (P0) C57BL/6 mice as described previously (43) with minor modifications. Briefly, the mice were decapitated, the brain structures were removed. Cortexes from 3 mice were dissected and rinsed in cold Hank’s balanced salt solution (HBSS), and the meninges were carefully stripped off. Tissue was triturated and, after centrifugation, the pellet was resuspended in astrocyte culture medium (DMEM containing 10% fetal bovine serum (Giboco), 0.5% Penicillin/Streptomycin and 1% GlutaMAX supplement (Giboco). Cells were plated in T-75 flasks (Corning) previously coated with poly-L-lysine (Sigma) and incubated at 37°C in a humidified 5% CO_2_, 95% air chamber for 7–8 days until reaching confluence. After growing to confluence, cells were shaken for 7 h at 240 rpm on an orbital shaker to remove microglia and oligodendrocyte precursor cells. For morphology analysis, purified astrocytes were plated onto poly-D-lysine-coated coverslips at 50,000 cells/well on a 12-well plate before the day of the transfection (at 9-13 *in vitro*). EGFP plasmids were then transfected into astrocytes using a modified Lipofectamine protocol with Lipofectamine 3000 (Invitrogen) according the manufacturer’s instructions and applied directly to cultured cells and allowing overnight (24 h) expression.

### Immunohistochemical (IHC) evaluations

IHC was performed as previously reported (41). For immunofluorescence in cultured astrocytes, cells were fixed with 4% paraformaldehyde (15 min, 0°C), rinsed with glycine solution (30 mM in 0.1 M PBS solution), and permeabilized with Triton X-100 (0.2%, 3 min). Preincubation was performed with 10% NGS for 1 h at 37°C. The following primary antibodies were used: mouse anti-NeuN (1:1000; Abcam #ab104224), mouse anti-GFAP (1:500, Millipore #MAB360), rabbit anti-GFAP (1:500, Proteintech #16825-1-AP), guinea pig anti-S100β (1:500, Synaptic Systems #287 004), rabbit anti-Ezrin (1:100, Cell Signaling #3145), mouse anti-vGluT1 (1:200, Millipore #MAB5502), rabbit ant-PSD95 (1:100, Invitrogen #51-6900). The following Alexa conjugated secondary antibodies were used: goat anti-mouse 488 (1:1000, Abcam #ab150113), goat rabbit-488 (1:1000, Abcam #ab150077), goat anti-rabbit 647 (1:500, Abcam #ab150079) and goat anti-guinea pig 488 (1:500, Abcam #ab150185). Immunofluorescence of Ezrin was performed by using an Alexa Fluor™ 488 Tyramide SuperBoost™ Kit (Invitrogen #B40912, and #B40941) according to the manufacturer’s protocols. After nucleus labeling with DAPI, the coverslips were mounted on slides using anti-fade solution. For quantification of fluorescent intensity, sections from the two groups were stained and imaged with exactly the same protocol.

#### Synaptic counting

After vGluT1 and PSD95 staining, high magnification 60× objective lens plus 4× optical zoom Z-stack images were obtained using the Nikon A1R^+^ multiphoton microscope. To obtain a high signal-to-noise ratio, ER function of Nikon A1R^+^multiphoton microscope was further used. Roughly half of each image area contained a mCherry positive astrocyte, and the remainder of the image was mCherry negative. 5 µm thick Z-stacks of 20 optical sections were imaged with all three channels: vGluT1 (green), mCherry (red), and PSD95 (magenta), and four consecutive optical sections (thickness of 1 μm) were then extracted and imported to Imaris for further analysis. The number of co-localized synaptic puncta of excitatory intracortical (vGluT1/PSD95) was obtained using the Imaris plugin Surface-surface colocalization after 3D reconstructing of vGluT1/PSD95 with *Surface* function. For each image, co-localized synaptic puncta were quantified in mCherry^+^ and mCherry^−^ domains of ROIs of 225 µm^3^ (15 µm × 15 µm × 1 µm) which were devoid of regions with neuronal cell bodies (areas lacking synaptic puncta). Synaptic puncta quantification was calculated as a density (synaptic puncta counts/ 225 µm^3^).

### Statistics

All statistical analyses were performed using Origin 9 software (OriginLab). Data were represented as means ± s.e.m; nonparametric data were represented as medians. Data fitting a parametric distribution were tested for significance using analysis of paired and unpaired Student’s two-tailed *t* tests; Data fitting a nonparametric distribution were tested for significance using two-tailed Mann–Whitney. Data with more than two groups were tested for significance using one-way ANOVA test. Significance was defined as *P <* 0.05.

## Acknowledgments

Thanks to Mr. XiaoWei Liu and Dr. Biao Dong for AAVs packaging. Thanks to Hong Kong Sun Joy Instrument Trading Co., Ltd for sharing confocal microscope and Imaris software. Thanks to Core Facility of West China Hospital, Public Health and Preventive Medicine Provincial Experiment Teaching Center at Sichuan University, Analysis and Testing Center of Sichuan University, Food Safety Monitoring and Risk Assessment Key Laboratory of Sichuan Province for sharing instruments. Thanks to ZEISS Microscopy Customer Center Shanghai for helping with some pilot experiments with confocal microscopy. We thank Prof. Baljit S Khakh, Prof. Ji Xu and Prof. Xiaoping Tong for critical reading of the manuscript.

## Supplementary information

**Supplementary Table 1. Artery blood gas analysis of P7 mice**

**Supplementary Fig 1. Early Sevo exposure neither affect astrocyte soma and primary branch morphology nor induce apparent astrogliosis in the somatosensory cortex**

(**A**) Representative confocal image and 3D reconstructed soma and primary branches of a cortical astrocyte. Scale bar, 10 μm. (**B**) **left**: quantification of astrocytic soma volume at P8 (*P* = 0.216, unpaired *t* test), P14 (*P* = 0.652, unpaired *t* test) and P21 (*P* = 0.449, unpaired *t* test); **middle**: quantification of primary branches volume of astrocytes in the somatosensory cortex at P8 (*P* = 0.098, unpaired *t* test), P14 (*P* = 0.826, unpaired *t* test) and P21 (*P* = 0.798, unpaired *t* test); **right**: average astrocytes number of primary branches in the somatosensory cortex at P8 (*P* = 0.592, Mann-Whitney test), P14 (*P* = 0.219, Mann-Whitney test) and P21 (*P* = 0.065, Mann-Whitney test). (**C**) Representative confocal images (left), zoom in (right top) and 3D reconstruction (right bottom) of GFAP in the somatosensory cortex at P14. Scale bars, 100 μm (left), 10 μm (right). (**D**) Quantification of GFAP volume (*P* = 0.263, unpaired *t* test); (**left**) and GFAP^+^ cells density (*P* = 0.731, unpaired *t* test) (**right**). n.s., not significant. Data are shown as mean ± s.e.m.

**Supplementary Fig 2. Early Lengthy Sevo exposure did not lead to astrocyte morphological deficits in the hippocampal CA1sr and DG-mo at P14**

(**A)** Diagram of hippocampal CA1sr and DG-mo. (**B**) Representative fluorescent images and distal fine processes reconstructions of astrocytes in the hippocampal CA1sr and DG-mo of P14 mice. Scale bars, 10 μm. (**C, D**) Average distal and proximal fine processes volume of astrocytes in the hippocampal CA1sr and DG-mo of P14 mice (CA1sr distal fine processes: *P* = 0.888; DG-mo distal fine processes: *P* = 0.278; CA1sr proximal fine processes: *P* = 0.009; DG-mo proximal fine processes: *P* = 0.046, unpaired *t* test). (**E, F, G, H**) Quantification of astrocytic territory (**E**) and soma (**F**) volume, primary branches volume (**G**) and number of primary branches (**H**), respectively, (CA1sr territory volume: *P* = 0.737; DG-mo territory volume: *P* = 0.20; CA1sr soma volume: *P* = 0.895; DG-mo soma volume: *P* = 0.347; CA1sr primary branches volume: *P* = 0.553; DG-mo number of primary branches: *P* = 0.797, unpaired *t* test; DG-mo primary branches volume: *P* = 0.037; CA1sr number of primary branches: *P* = 0.905, Mann-Whitney test). \**P* < 0.05, \*\**P* < 0.01, n.s., not significant. Data are shown as mean ± s.e.m.

**Supplementary Fig 3. 1 h Sevo exposure to P7 mice or 4 h exposure to P42-50 mice did not impair astrocyte morphogenesis**

(**A**) Experiment protocol for Sevo exposure and morphological assessment. (**B)** Fluorescent images and reconstructed distal fine processes of astrocytes. Scale bars, 10 μm. (**C, D**) Quantification of astrocytic distal and proximal fine processes volume. (P42-50 distal fine processes: *P* = 0.30, unpaired *t* test; P14 distal fine processes: *P* = 0.486, Mann-Whitney test; P42-50 proximal fine processes: *P* = 0.915; P14 proximal fine processes: *P* = 0.552, unpaired *t* test). (**E, F, G, H)** Quantification of astrocytic territory **(E)** and soma **(F)** volume, volume **(G)** and number **(H)** of primary branches, respectively (P42-50 soma volume: *P* = 0.173; P14 soma volume: *P* = 0.391; P42-50 territory volume: *P* = 0.268; P14 territory volume: *P* = 0.439, unpaired *t* test; P42-50 primary branches volume: *P* = 0.675; P14 primary branches volume: *P* = 0.187; P42-50 number of primary branches: *P* = 0.978; P14 number of primary branches: *P* = 0.181, Mann-Whitney test). n.s., not significant. Data are shown as mean ± s.e.m.

**Supplementary Fig 4. Unaltered total and mushroom apical dendritic spine density in the hippocampal CA1sr at P21**

**(A)** Confocal images of apical dendritic spines in the hippocampal CA1sr. Scale bar, 2 μm. **(B)** Quantification of total (*P* = 0.079, unpaired *t* test) and mushroom (*P* = 0.267, unpaired *t* test) apical dendritic spine density. n.s., not significant. Data are shown as mean ± s.e.m.

**Supplementary Fig 5. eEPSCs pharmacological blockade and kinetics in cortical pyramidal neurons from Control and Sevo group mice**

(**A**) Traces depicting pharmacological eEPSCs at −60 mV and +40 mV. (**B, C**) Quantification of the decay kinetics (weighted time constants) of AMPAR-mediated eEPSCs and NMDAR-mediated eEPSCs in Control and Sevo group mice (**B**) *P* = 0.167, Mann-Whitney test; (**C**) *P* = 0.119, Mann-Whitney test). \*\**P* < 0.01, n.s., not significant. Data are shown as mean ± s.e.m.

**Supplementary Fig 6. Stereotactic injection of AAV5⋅gfaABC_1_D⋅GCaMP6f into the cortex of P0 mice did not induce apparent astrogliosis at P14.** Fluorescent images of auto-EGFP (green) and GFAP (red) in the cortex of WT and GCaMP6f injected mice at P14. Scale bars, 200 μm.

**Supplementary Fig 7. Inhibition of basal Ca^2+^ level and spontaneous Ca^2+^ signals following acute Sevo exposure**

(**A**) Quantification of acute Sevo application on cortical astrocyte basal Ca^2+^ from adult mice at P56 (*P* = 0.004, unpaired *t* test). (**B**) Quantification of the two types of spontaneous Ca^2+^ signals properties including frequency, peak amplitude and half-width before and after acute Sevo exposure (*P* = 0.009, paired *t* test, for waves frequency; *P* = 0.997, paired *t* test for microdomains frequency; *P* = 0.009, Mann-Whitney test for waves peak amplitude; *P* = 0.188, Mann-Whitney test for microdomains peak amplitude; *P* = 0.717, Mann-Whitney test for waves half-width; *P* = 0.742, Mann-Whitney test for microdomains half-width). n.s., not significant. Data are shown as mean ± s.e.m.

**Supplementary Fig 8. Unchanged Ezrin expression levels in the hippocampal CA1sr and DG-mo in Sevo group mice at P14**

(**A**) Representative fluorescent images in the hippocampus of Control and Sevo group mice at P14. Scale bars, 200 μm. (**B**) Quantification of Ezrin fluorescent intensity in the hippocampal CA1sr (n = 96 ROIs from 6 mice in Control group, n = 86 ROIs from 6 mice in Sevo group; *P* = 0.295, Mann-Whitney test) and DG-mo (n = 84 ROIs from 6 mice in Control group, n = 76 ROIs from 6 mice in Sevo group; *P* = 0.164, Mann-Whitney test). n.s., not significant. Data are shown as mean ± s.e.m.

**Supplementary Fig 9. Ezrin-miRNAi, with high knock-down efficiency and cell-type specificity, lead to increased astrocyte soma volume but unchanged primary branches morphology in the cortex at P21**

(**A**) Fluorescent images of Ezrin and mCherry in the cortex of NC-miRNAi and Ezrin-miRNAi mice at P21. Scale bars, 100 μm. (**B**) Quantification of fold change in Ezrin fluorescent intensity, measured as (F_mCherry+_-F_mCherry-_)/ F_mCherry-_ (*P* = 0, Mann-Whitney test). (**C**) Schematic fluorescent images of S100β, mCherry and NeuN in the cortex of Ezrin-miRNAi injected mice at P21. Scale bars, 50 μm. (**D**) Quantification of mCherry^+^ cells colocalized with S100β and NeuN. (**E**) Images of GFAP and mCherry in the cortex of Ezrin-miRNAi and NC-miRNAi injected mice at P21. Scale bars, 20 μm. (**F**) Confocal image with 3D reconstructed soma and primary branches of mCherry labeled astrocyte. Scale bar, 10 μm. (**G**) Quantification of astrocyte soma volume (**left**), primary branches volume (**middle**) and number of primary branches (**right**), respectively, in NC-miRNAi and Ezrin-miRNAi injected mice (soma volume: *P* = 6.4E-4, number of primary branches: *P* = 0.095, Mann-Whitney test; primary branches volume: *P* = 0.072, unpaired *t* test). \*\*\**P* < 0.001, n.s., not significant. Data are shown as mean ± s.e.m.

## Abbreviations

GAs: general anesthetics
Sevo: sevoflurance
CA1sr: CA1 stratum radiatum
DG-mo: molecular layer of dentate gyrus
SBF-SEM: serial block face scanning electron microscopy
GFAP: glial fibrillary acidic protein
GLAST: glutamate-aspartate transporter
eEPSCs: evoked EPSCs
mEPSCs: miniature EPSCs
MACS: magnetic cell sorting
RT-qPCR: quantitative reverse transcription PCR
GCaMP6f: AAV5•gfaABC1D•GCaMP6f
Ezrin-miRNAi: AAV8•gfaABC1D•mCherry•Ezrin-miRNAi
NC-miRNAi: AAV8•gfaABC1D•mCherry•NC-miRNAi

## References

1. Vutskits L, Xie Z. Lasting impact of general anaesthesia on the brain: mechanisms and relevance. Nat Rev Neurosci. 2016;17(11):705–17.

2. Soriano SG, Vutskits L, Jevtovic-Todorovic V, Hemmings HC, Neurotoxicology BJA, Neuroplasticity Study G. Thinking, fast and slow: highlights from the 2016 BJA seminar on anaesthetic neurotoxicity and neuroplasticity. Br J Anaesth. 2017;119(3):443–7.

3. Warner DO, Zaccariello MJ, Katusic SK, Schroeder DR, Hanson AC, Schulte PJ, et al. Neuropsychological and Behavioral Outcomes after Exposure of Young Children to Procedures Requiring General Anesthesia: The Mayo Anesthesia Safety in Kids (MASK) Study. Anesthesiology. 2018;129(1):89–105.

4. Hu D, Flick RP, Zaccariello MJ, Colligan RC, Katusic SK, Schroeder DR, et al. Association between Exposure of Young Children to Procedures Requiring General Anesthesia and Learning and Behavioral Outcomes in a Population-based Birth Cohort. Anesthesiology. 2017;127(2):227–40.

5. Pavkovic Z, Milanovic D, Ruzdijic S, Kanazir S, Pesic V. The influence of propofol anesthesia exposure on nonaversive memory retrieval and expression of molecules involved in memory process in the dorsal hippocampus in peripubertal rats. Paediatr Anaesth. 2018;28(6):537–46.

6. Satomoto M, Satoh Y, Terui K, Miyao H, Takishima K, Ito M, et al. Neonatal exposure to sevoflurane induces abnormal social behaviors and deficits in fear conditioning in mice. Anesthesiology. 2009;110(3):628–37.

7. Zhong L, Luo F, Zhao W, Feng Y, Wu L, Lin J, et al. Propofol exposure during late stages of pregnancy impairs learning and memory in rat offspring via the BDNF-TrkB signalling pathway. J Cell Mol Med. 2016;20(10):1920–31.

8. Jevtovic-Todorovic V, Hartman RE, Izumi Y, Benshoff ND, Dikranian K, Zorumski CF, et al. Early exposure to common anesthetic agents causes widespread neurodegeneration in the developing rat brain and persistent learning deficits. J Neurosci. 2003;23(3):876–82.

9. Allen NJ, Eroglu C. Cell Biology of Astrocyte-Synapse Interactions. Neuron. 2017;96(3):697–708.

10. Farhy-Tselnicker I, Allen NJ. Astrocytes, neurons, synapses: a tripartite view on cortical circuit development. Neural Dev. 2018;13(1):7.

11. Wang W, Lu R, Feng DY, Zhang H. Sevoflurane Inhibits Glutamate-Aspartate Transporter and Glial Fibrillary Acidic Protein Expression in Hippocampal Astrocytes of Neonatal Rats Through the Janus Kinase/Signal Transducer and Activator of Transcription (JAK/STAT) Pathway. Anesth Analg. 2016;123(1):93–102.

12. Liu Y, Yan Y, Inagaki Y, Logan S, Bosnjak ZJ, Bai X. Insufficient Astrocyte-Derived Brain-Derived Neurotrophic Factor Contributes to Propofol-Induced Neuron Death Through Akt/Glycogen Synthase Kinase 3beta/Mitochondrial Fission Pathway. Anesth Analg. 2017;125(1):241–54.

13. Ryu YK, Khan S, Smith SC, Mintz CD. Isoflurane impairs the capacity of astrocytes to support neuronal development in a mouse dissociated coculture model. J Neurosurg Anesthesiol. 2014;26(4):363–8.

14. Wilder RT, Flick RP, Sprung J, Katusic SK, Barbaresi WJ, Mickelson C, et al. Early exposure to anesthesia and learning disabilities in a population-based birth cohort. Anesthesiology. 2009;110(4):796–804.

15. Octeau JC, Chai H, Jiang R, Bonanno SL, Martin KC, Khakh BS. An Optical Neuron-Astrocyte Proximity Assay at Synaptic Distance Scales. Neuron. 2018;98(1):49–66.

16. Araque A, Parpura V, Sanzgiri RP, Haydon PG. Tripartite synapses: glia, the unacknowledged partner. Trends Neurosci. 1999;22(5):208–15.

17. Thrane AS, Rangroo Thrane V, Zeppenfeld D, Lou N, Xu Q, Nagelhus EA, et al. General anesthesia selectively disrupts astrocyte calcium signaling in the awake mouse cortex. Proc Natl Acad Sci U S A. 2012;109(46):18974–9.

18. Franks NP, Lieb WR. Temperature dependence of the potency of volatile general anesthetics: implications for in vitro experiments. Anesthesiology. 1996;84(3):716–20.

19. Wang Y, Shi G, Miller DJ, Wang Y, Wang C, Broussard G, et al. Automated Functional Analysis of Astrocytes from Chronic Time-Lapse Calcium Imaging Data. Front Neuroinform. 2017(11):48.

20. Srinivasan R, Huang BS, Venugopal S, Johnston AD, Chai H, Zeng H, et al. Ca(2+) signaling in astrocytes from Ip3r2(−/−) mice in brain slices and during startle responses in vivo. Nat Neurosci. 2015;18(5):708–17.

21. Derouiche A, Frotscher M. Peripheral astrocyte processes: monitoring by selective immunostaining for the actin-binding ERM proteins. Glia. 2001;36(3):330–41.

22. Lavialle M, Aumann G, Anlauf E, Prols F, Arpin M, Derouiche A. Structural plasticity of perisynaptic astrocyte processes involves ezrin and metabotropic glutamate receptors. Proc Natl Acad Sci U S A. 2011;108(31):12915–9.

23. Briner A, Nikonenko I, De Roo M, Dayer A, Muller D, Vutskits L. Developmental Stage-dependent persistent impact of propofol anesthesia on dendritic spines in the rat medial prefrontal cortex. Anesthesiology. 2011;115(2):282–93.

24. Xiao H, Liu B, Chen Y, Zhang J. Learning, memory and synaptic plasticity in hippocampus in rats exposed to sevoflurane. Int J Dev Neurosci. 2016;48:38–49.

25. Amrock LG, Starner ML, Murphy KL, Baxter MG. Long-term effects of single or multiple neonatal sevoflurane exposures on rat hippocampal ultrastructure. Anesthesiology. 2015;122(1):87–95.

26. Kang E, Jiang D, Ryu YK, Lim S, Kwak M, Gray CD, et al. Early postnatal exposure to isoflurane causes cognitive deficits and disrupts development of newborn hippocampal neurons via activation of the mTOR pathway. PLoS Biol. 2017;15(7):e2001246.

27. Oliveira JF, Sardinha VM, Guerra-Gomes S, Araque A, Sousa N. Do stars govern our actions? Astrocyte involvement in rodent behavior. Trends in neurosciences. 2015;38(9):535–49.

28. Shigetomi E, Patel S, Khakh BS. Probing the Complexities of Astrocyte Calcium Signaling. Trends Cell Biol. 2016;26(4):300–12.

29. Zheng K, Bard L, Reynolds JP, King C, Jensen TP, Gourine AV, et al. Time-Resolved Imaging Reveals Heterogeneous Landscapes of Nanomolar Ca(2+) in Neurons and Astroglia. Neuron. 2015;88(2):277–88.

30. Campagna JA, Miller KW, Forman SA. Mechanisms of actions of inhaled anesthetics. N Engl J Med. 2003;348(21):2110–24.

31. Bernardinelli Y, Randall J, Janett E, Nikonenko I, Konig S, Jones EV, et al. Activity-dependent structural plasticity of perisynaptic astrocytic domains promotes excitatory synapse stability. Curr Biol. 2014;24(15):1679–88.

32. Hanse E, Seth H, Riebe I. AMPA-silent synapses in brain development and pathology. Nat Rev Neurosci. 2013;14(12):839–50.

33. Chung WS, Allen NJ, Eroglu C. Astrocytes Control Synapse Formation, Function, and Elimination. Cold Spring Harb Perspect Biol. 2015;7(9):a020370.

34. Stogsdill JA, Ramirez J, Liu D, Kim YH, Baldwin KT, Enustun E, et al. Astrocytic neuroligins control astrocyte morphogenesis and synaptogenesis. Nature. 2017;551(7679):192–7.

35. Femenia T, Gimenez-Cassina A, Codeluppi S, Fernandez-Zafra T, Katsu-Jimenez Y, Terrando N, et al. Disrupted Neuroglial Metabolic Coupling after Peripheral Surgery. J Neurosci. 2018;38(2):452–64.

36. Tong X, Ao Y, Faas GC, Nwaobi SE, Xu J, Haustein MD, et al. Astrocyte Kir4.1 ion channel deficits contribute to neuronal dysfunction in Huntington’s disease model mice. Nat Neurosci. 2014;17(5):694–70.

37. Ballesteros-Yanez I, Benavides-Piccione R, Bourgeois JP, Changeux JP, DeFelipe J. Alterations of cortical pyramidal neurons in mice lacking high-affinity nicotinic receptors. Proc Natl Acad Sci U S A. 2010;107(25):11567–72.

38. Li H, Li Y, Lei Z, Wang K, Guo A. Transformation of odor selectivity from projection neurons to single mushroom body neurons mapped with dual-color calcium imaging. Proc Natl Acad Sci U S A. 2013;110(29):12084–9.

39. Medvedev N, Popov V, Henneberger C, Kraev I, Rusakov DA, Stewart MG. Glia selectively approach synapses on thin dendritic spines. Philos Trans R Soc Lond B Biol Sci. 2014;369(1654):20140047.

40. Kim JY, Grunke SD, Levites Y, Golde TE, Jankowsky JL. Intracerebroventricular viral injection of the neonatal mouse brain for persistent and widespread neuronal transduction. J Vis Exp. 2014(91):51863.

41. Jiang R, Diaz-Castro B, Looger LL, Khakh BS. Dysfunctional Calcium and Glutamate Signaling in Striatal Astrocytes from Huntington’s Disease Model Mice. J Neurosci. 2016;36(12):3453–70.

42. Foo LC. Purification of astrocytes from transgenic rodents by fluorescence-activated cell sorting. Cold Spring Harb Protoc. 2013;2013(6):551–60.

43. Schildge S, Bohrer C, Beck K, Schachtrup C. Isolation and culture of mouse cortical astrocytes. J Vis Exp. 2013;(71).(pii):50079.

